# The MRE11-Rad50-Nbs1 complex acts on 3’ DNA-protein covalent complexes

**DOI:** 10.64898/2026.11.23.768626

**Authors:** Yang Ye, Sanjay Kumar, Bokun Yin, Jingwei Xue, Nicolette A. Ender, Hannah K. Barnes, Omar Escalante, Liton K. Saha, Hiroyuki Sasanuma, Shunichi Takeda, Tanya T. Paull

## Abstract

DNA-protein crosslinks (DPCs) are complex lesions that block replication, transcription, and other DNA transactions. Here we investigate repair of 3’ DPC lesions using the site-specific Flp tyrosine recombinase H305L mutant that forms stable covalent complexes. In human cells, removal or catalytic inhibition of MRE11 stabilizes Flp DPCs in parallel to a proteasome-dependent pathway. High levels of Flp DPCs are also observed in budding yeast, where MRE11 deletion or mutation of the nuclease domain increases covalent Flp DPC accumulation. Mass spectrometry-based identification of endogenous DPC factors in human cells identifies TOP1 covalent complexes as the primary 3’ lesion that is stabilized in the absence of MRE11 nuclease function, in addition to 5’ DPCs generated by TOP2 family enzymes. This work characterizes an important role for the MRE11-RAD50-NBS1 complex in processing of 3’ lesions, in addition to its well-established functions at 5’ DPCs and in 5’ strand resection during homologous recombination.

## Introduction

DNA–protein crosslinks (DPCs) are bulky DNA lesions in which proteins become covalently linked to DNA, creating physical barriers to DNA replication, transcription, and other DNA transactions ^1^. DPCs can arise as trapped intermediates of DNA-processing enzymes, including topoisomerases, or from endogenous and environmental crosslinking agents such as aldehydes. Because DPCs vary widely in protein size, linkage chemistry, and surrounding DNA structure, cells use multiple mechanisms to remove them ^1–3^. These include proteolytic pathways involving the proteasome and DPC-specific proteases such as SPRTN, direct hydrolysis of phosphotyrosyl linkages by tyrosyl-DNA phosphodiesterases, and nuclease-mediated excision of protein-linked DNA. The relative contribution of these pathways depends strongly on the identity and structure of the lesion, suggesting that DPC repair is organized as a network of partially overlapping mechanisms rather than a single universal pathway.

The MRE11–RAD50–NBS1 (MRN) complex is well established as a central regulator of DNA double-strand break repair and also has important functions in processing protein-blocked DNA ends ^4–7^. In budding yeast, the related MRX complex and Sae2 promote removal of 5′ Spo11-linked meiotic DNA ends, and biochemical studies have shown that MRX–Sae2 and human MRN can introduce endonucleolytic incisions adjacent to protein-blocked DNA ends and subsequently process the resulting DNA intermediates ^4,5^. Cellular studies have further implicated MRE11 and CtIP in removal of TOP2 cleavage complexes, which contain 5′-phosphotyrosyl protein-DNA linkages ^7,8^. MRE11 has also been linked genetically and biochemically to repair of TOP1-associated 3′-phosphotyrosyl lesions, although the requirement for MRE11 nuclease activity is strongly influenced by DNA-end architecture and lesion context ^6,9,10^. Despite these findings, the role of MRE11 in processing a defined 3′-protein adduct in cells remains poorly resolved. Moreover, many cellular assays infer DPC processing from downstream outcomes such as resection, homologous recombination, chromosome breakage, or cell survival. Because MRN also functions directly in these downstream DNA repair processes, such measurements do not necessarily distinguish removal of the protein adduct itself from subsequent repair of the associated DNA lesion.

A major challenge in defining individual DPC repair mechanisms is that different experimental systems capture different aspects of DPC biology. Broad-spectrum crosslinking agents such as formaldehyde generate heterogeneous populations of DPCs involving many proteins, linkage chemistries, and genomic contexts, whereas topoisomerase poisons such as camptothecin and etoposide generate biologically relevant, chemically defined TOP1- and TOP2-DNA cleavage complexes across the genome ^1,3,11^. These approaches have been essential for identifying DPC repair pathways and for understanding the consequences of endogenous protein-DNA adducts, but their genome-wide distribution and dependence on endogenous enzyme function can make it difficult to isolate the processing of a single defined lesion from downstream effects on replication, transcription, and DNA repair. A complementary approach is therefore to use a genetically defined DPC that can be induced at a predetermined genomic locus and monitored directly.

To investigate DPC processing, we employed the site-specific tyrosine recombinase Flp, originally from budding yeast. During recombination, Flp forms a transient covalent intermediate with a 3′ DNA strand at its target site (FRT), which is subsequently resolved by nucleophilic attack by the recombining DNA molecule ^12^. The H305L mutation in Flp traps this intermediate by impairing subsequent strand joining ^12,13^. This property has been exploited in the Flp-nick system to study the consequences of a site-specific protein-bound nick in yeast and, more recently, mammalian cells ^14–19^. The FlpH305L–FRT system thus provides a site-specific DPC model in which the persistence of a defined 3′-phosphotyrosyl covalent protein linkage can be measured directly at a known genomic site.

Here, we established inducible FlpH305L–FRT systems in mammalian cells and budding yeast and used them to directly monitor the persistence of a defined 3′-phosphotyrosyl DPC. A focused genetic and pharmacological analysis identified multiple pathways contributing to Flp-DPC processing, including TDP1 and the structure-selective nucleases MUS81 and XPF. Most notably, depletion or catalytic inhibition of MRE11 increased Flp-DPC persistence in mammalian cells, while deletion of MRE11, mutation of MRE11 catalytic residues, and loss of Sae2 produced strong DPC accumulation in budding yeast. MRE11 was also enriched at the induced Flp lesion, supporting a direct role in DPC processing. Finally, perturbation of MRE11 increased the recovery of endogenous DNA-associated topoisomerases and increased TOP1 cleavage complexes, extending the relevance of these findings beyond the engineered Flp substrate. Together, these results identify MRE11 nuclease activity as a conserved component of 3′-protein adduct processing and establish the FlpH305L–FRT system as a tractable model for dissecting DPC repair mechanisms in eukaryotic cells.

## Results

### Establishment and validation of an inducible site-specific FlpHL–FRT DPC system

We used the H305L mutant of Flp recombinase (FlpHL) to generate a site-specific DNA–protein crosslink at a stably integrated FRT site. Flp cleaves DNA at two sites within the FRT sequence and forms a covalent intermediate between its catalytic tyrosine and the 3′ DNA end. The H305L mutation blocks resolution of this intermediate, resulting in formation of a FlpHL–FRT DPC (Fig. 1A). To control DPC formation, FLAG-tagged FlpHL was expressed from a doxycycline-inducible promoter. Doxycycline treatment induced FLAG-FlpHL expression in HEK-293 and U2OS cells, with increasing expression over the 8-h induction period in HEK-293 cells (Fig. 1B).

**Figure 1.**
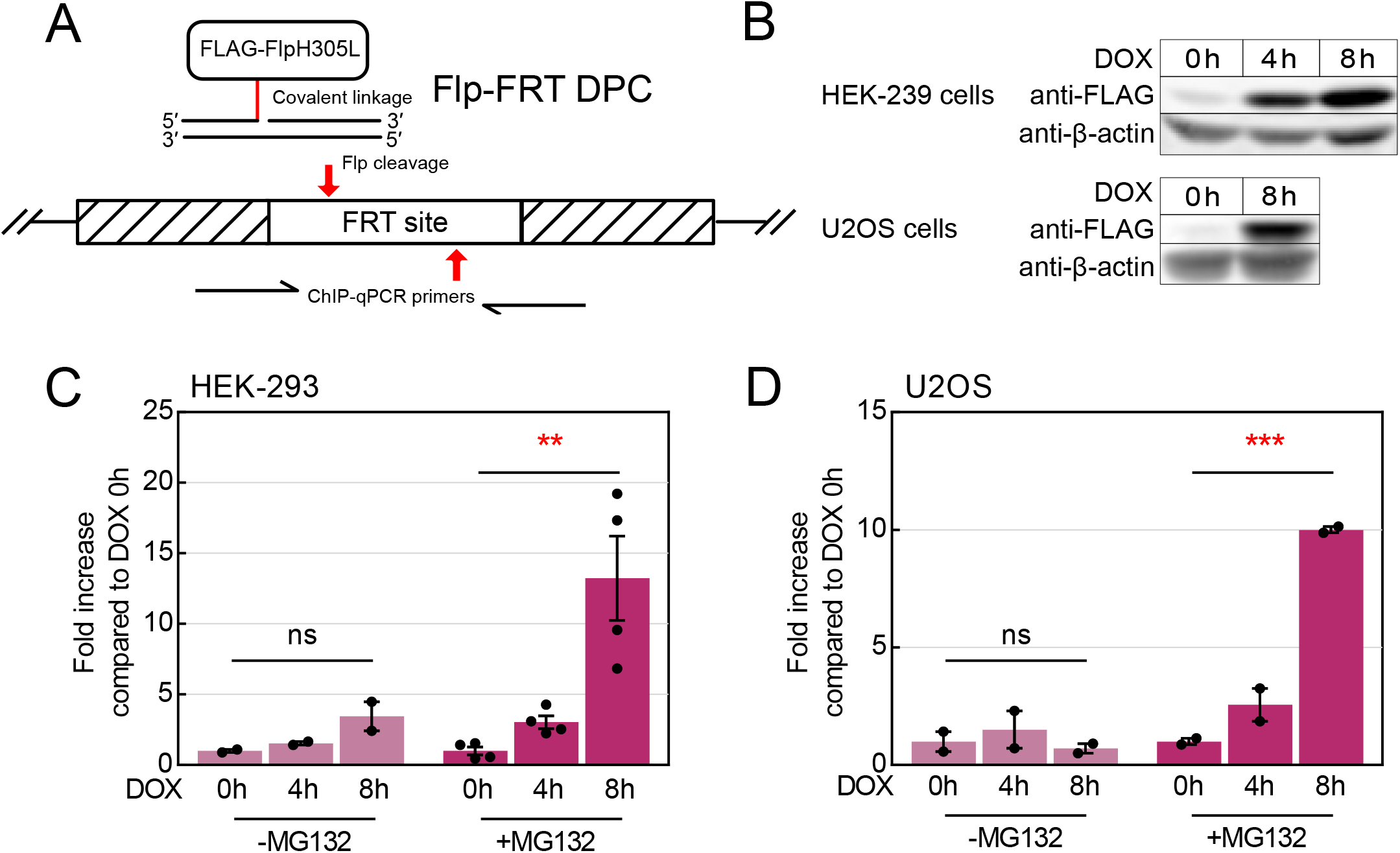
Establishment and validation of an inducible site-specific FlpHL–FRT DPC system. (A) Schematic of the FlpHL–FRT DPC system. FlpH305L (FlpHL) cleaves DNA at the FRT site and forms a covalent linkage with the 3′ DNA end on one or both strands. The H305L mutation blocks resolution of this intermediate, resulting in formation of a FlpHL–FRT DNA–protein crosslink (DPC). The FRT site contains two Flp cleavage sites flanking a central spacer region; one or both sites can form a DPC. The primer pair used for ChIP-qPCR analysis of the integrated FRT locus in HEK-293 and U2OS Flp-In T-REx cells is indicated below the schematic. (B) Western blot analysis of doxycycline-induced FLAG-FlpHL expression in HEK-293 and U2OS cells. Cells were treated with doxycycline (DOX; 1 μg/mL) for the indicated times, and FLAG-FlpHL expression was detected using an anti-FLAG antibody. β-actin was used as a loading control. (C, D) ChIP-qPCR analysis of FlpHL–FRT DPC accumulation in HEK-293 (C) and U2OS (D) Flp-In T-REx cells following doxycycline induction in the presence or absence of MG132. Cells were treated with DOX (1 μg/mL) and, where indicated, MG132 (30 μM) for the indicated times. FLAG-FlpHL associated with the integrated FRT site was measured by anti-FLAG ChIP-qPCR. Values were normalized to the corresponding 0-h sample for each treatment condition. Data represent 2–4 biological replicates for C and two biological replicates for D. Individual points represent biological replicates; bars show mean ± SEM. For C and D, statistical significance was determined using a two-tailed Student’s *t* test assuming equal variance. Comparisons are indicated by horizontal lines. ** and *** indicate *P* < 0.01 and *P* < 0.001, respectively; ns, not significant.

We next asked whether FlpHL–FRT DPCs could be detected at the integrated FRT locus by performing ChIP in the absence of formaldehyde crosslinking followed by qPCR quantitation. Experiments were first performed in HEK-293 and U2OS Flp-In T-REx cells carrying a single integrated FRT site in the genome. FLAG-FlpHL associated with the FRT locus was measured by anti-FLAG ChIP followed by qPCR using primers spanning the integration site. With these primers, a single strand FLAG-FlpHL is recovered (either top or bottom strand, top strand DPC depicted in Fig. 1A). In the absence of the proteasome inhibitor MG132, only limited accumulation of FLAG-FlpHL was detected following doxycycline induction in either cell line. In contrast, MG132 treatment increased FLAG-FlpHL enrichment at the FRT locus in both cell lines, with the highest levels observed after 8 h of doxycycline treatment (Fig. 1, C and D). These results show that FlpHL–FRT DPC accumulation can be monitored by anti-FLAG ChIP-qPCR in human cells and that proteasome inhibition increases the steady-state level of FLAG-FlpHL associated with the FRT locus.

### A focused genetic and pharmacological screen identifies factors contributing to FlpHL–FRT DPC processing

Having established the FlpHL–FRT DPC assay, we next adapted the system to the human lymphoblast cell line TK6 to take advantage of a panel of defined DNA repair-deficient backgrounds ^7,20–24^. An FRT site was introduced within a heterologous transgene using a stably integrated lentiviral construct in addition to a lentiviral construct expressing doxycycline-inducible FLAG-FlpHL. Three ChIP-qPCR primer sets were designed to detect regions upstream of, across, and downstream of the FRT site (Fig. 2A). Doxycycline treatment induced FLAG-FlpHL expression in TK6 cells within 8 h, similar to our results in HEK-293 and U2OS cells (Fig. 2B). In wild-type cells, however, MG132 produced only a modest increase in FLAG-FlpHL enrichment compared with cells treated without MG132, reaching statistical significance only at the downstream region (Fig. 2C).

**Figure 2.**
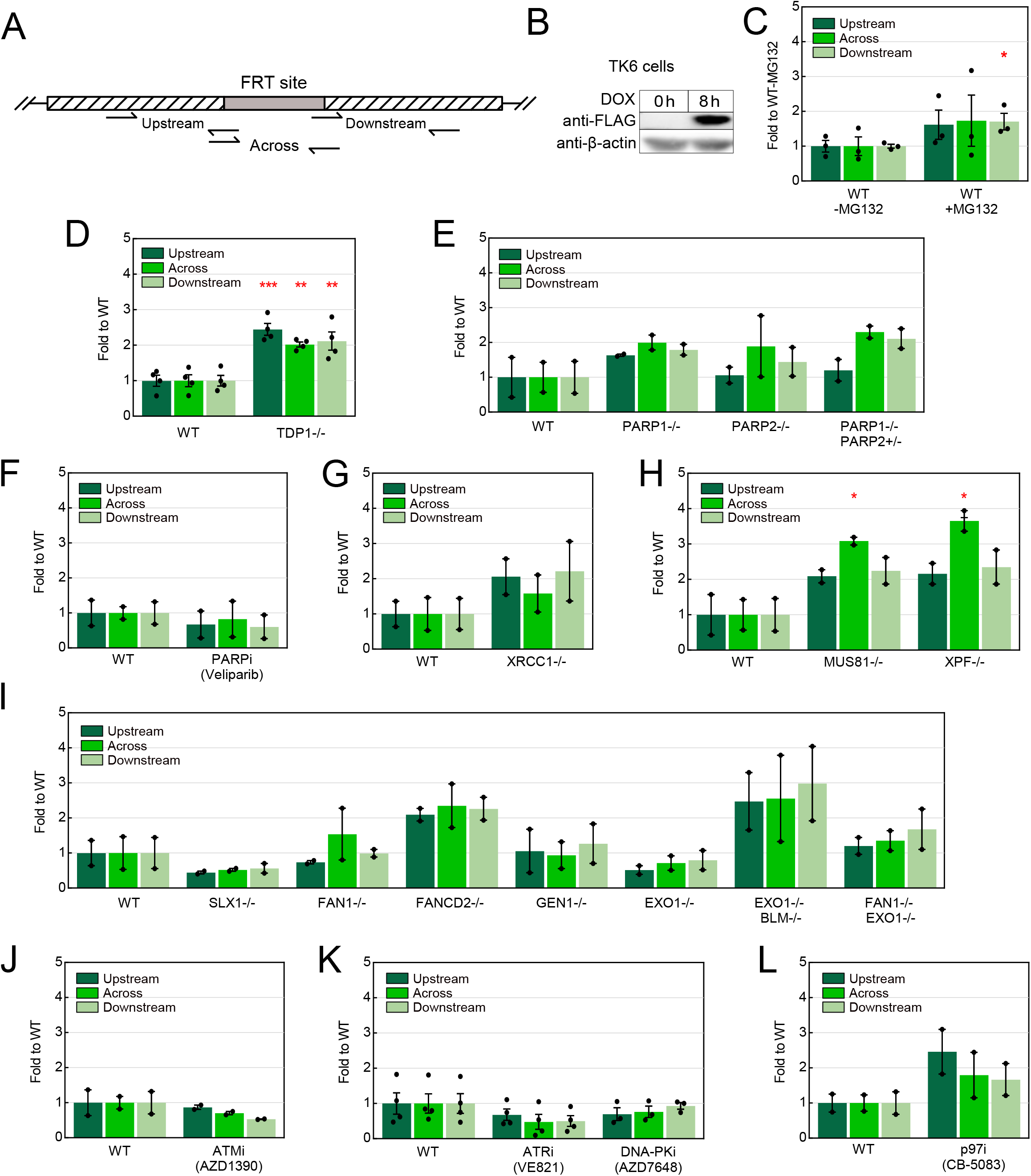
A focused genetic and pharmacological screen identifies factors contributing to FlpHL–FRT DPC processing. (A) Schematic of the engineered FRT locus and ChIP-qPCR primer design. The FRT site is integrated within a heterologous transgene. Three primer sets were used to measure FLAG-FlpHL enrichment upstream of, across, and downstream of the FRT site. (B) Western blot analysis of doxycycline-induced FLAG-FlpHL expression in TK6 cells carrying the engineered FRT locus. Cells were treated with DOX (1 μg/mL) for the indicated times, and FLAG-FlpHL expression was detected using an anti-FLAG antibody. β-actin was used as a loading control. (C) ChIP-qPCR analysis of FlpHL–FRT DPC accumulation in WT TK6 cells in the presence or absence of MG132. Cells carrying the FRT locus and doxycycline-inducible FLAG-FlpHL were treated with DOX for 8 h, with or without MG132 (30 μM). FLAG-FlpHL enrichment at the upstream, across, and downstream regions was measured by anti-FLAG ChIP-qPCR. Values for each primer set were normalized to the corresponding WT −MG132 value. (D) ChIP-qPCR analysis of FlpHL–FRT DPC accumulation in WT and *TDP1−/−* TK6 cells. Cells carrying the FRT locus and doxycycline-inducible FLAG-FlpHL were treated with DOX and MG132 (30 μM) for 8 h. FLAG-FlpHL enrichment at the upstream, across, and downstream regions was measured by anti-FLAG ChIP-qPCR. Values for each primer set were normalized to the corresponding WT value. (E) ChIP-qPCR analysis of FlpHL–FRT DPC accumulation in WT, PARP1−/−, PARP2−/−, and PARP1−/− PARP2+/− TK6 cells as in (D). (F) ChIP-qPCR analysis of FlpHL–FRT DPC accumulation in WT TK6 cells treated with the PARP inhibitor veliparib (10 μM). Cells were pretreated with veliparib overnight, followed by 8 h of DOX and MG132 treatment in the continued presence of veliparib. FLAG-FlpHL enrichment at the upstream, across, and downstream regions was measured by anti-FLAG ChIP-qPCR and normalized to the corresponding untreated WT value. (G) ChIP-qPCR analysis of FlpHL–FRT DPC accumulation in WT and XRCC1−/− TK6 cells. Cells were treated and analyzed as in D. (H) ChIP-qPCR analysis of FlpHL–FRT DPC accumulation in WT, *MUS81−/−*, and *XPF−/−* TK6 cells. Cells were treated and analyzed as in D. (I) ChIP-qPCR analysis of FlpHL–FRT DPC accumulation in WT and the indicated DNA repair-deficient TK6 cells, including *SLX1−/−, FAN1−/−, FANCD2−/−, GEN1−/−, EXO1−/−, EXO1−/− BLM−/−*, and *FAN1−/− EXO1−/−* cells. Cells were treated and analyzed as in D. (J) ChIP-qPCR analysis of FlpHL–FRT DPC accumulation in WT TK6 cells treated with the ATM inhibitor AZD1390 (1 μM). Cells were pretreated with AZD1390 overnight, followed by 8 h of DOX and MG132 treatment in the continued presence of the inhibitor. FLAG-FlpHL enrichment at the upstream, across, and downstream regions was measured by anti-FLAG ChIP-qPCR and normalized to the corresponding untreated WT value. (K) ChIP-qPCR analysis of FlpHL–FRT DPC accumulation in WT TK6 cells treated with the ATR inhibitor VE821 (1 μM) or the DNA-PK inhibitor AZD7648 (1 μM). Inhibitors were added for 8 h together with DOX and MG132. FLAG-FlpHL enrichment at the three regions was measured by anti-FLAG ChIP-qPCR and normalized to the corresponding untreated WT value. (L) ChIP-qPCR analysis of FlpHL–FRT DPC accumulation in WT TK6 cells treated with the p97 inhibitor CB-5083 (2.5 μM) for 8 h together with DOX and MG132. FLAG-FlpHL enrichment at the three regions was measured by anti-FLAG ChIP-qPCR and normalized to the corresponding untreated WT value. For C–L, data represent 2–4 biological replicates. Individual points represent biological replicates; bars show mean ± SEM. Unless otherwise indicated by horizontal comparison lines, statistical significance was determined by comparing each genotype or treatment with the corresponding WT control for the same primer set using a two-tailed Student’s *t* test assuming equal variance. *, **, and *** indicate *P* < 0.05, *P* < 0.01, and *P* < 0.001, respectively.

We first reasoned that TDP1, a tyrosyl-DNA phosphodiesterase involved in the removal of TOP1–DNA covalent intermediates, was a likely candidate for processing Flp adducts in TK6 cells because FlpHL forms a 3′-phosphotyrosyl linkage with DNA, similar to TOP1. Loss of TDP1 resulted in an approximately twofold increase in FLAG-FlpHL enrichment relative to wild-type cells with all three qPCR primer sets (Fig. 2D), consistent with a role for TDP1 in processing FlpHL–FRT DPCs. Similar accumulation of FlpHL DPC with the “across”, upstream, and downstream primer sets indicates that the DPCs formed in this system likely have Flp protein attached to only one strand.

Because asymmetric FlpHL cleavage generates a protein-blocked DNA strand and a single-strand break, we next examined factors involved in single-strand break repair. Loss of PARP1 or PARP2, either individually or in the *PARP1−/− PARP2+/−* background, produced only modest changes in FlpHL–FRT DPC levels compared with wild-type cells (Fig. 2E). Consistent with these results, treatment of wild-type TK6 cells with the PARP inhibitor veliparib did not increase DPC accumulation (Fig. 2F). XRCC1 deficiency also produced only a limited increase in FLAG-FlpHL enrichment (Fig. 2G). Thus, single-strand break repair factors have relatively little effect on steady-state FlpHL–FRT DPC levels under the conditions examined.

We next examined structure-selective nucleases and other DNA processing factors. Loss of MUS81 or XPF increased FLAG-FlpHL enrichment at the FRT locus, with the largest increases observed using the across primer set (Fig. 2H). In contrast, the remaining repair-deficient backgrounds generally showed smaller or more variable changes (Fig. 2I). FANCD2-deficient cells showed an approximately twofold increase in FLAG-FlpHL enrichment, whereas FAN1 deficiency did not produce a comparable effect. Similarly, loss of SLX1, GEN1, or EXO1 did not increase DPC accumulation. Combined loss of EXO1 and BLM resulted in higher FLAG-FlpHL enrichment than EXO1 deficiency alone, although the effect was variable, while combined loss of FAN1 and EXO1 produced only a modest increase. These results suggest that MUS81 and XPF could contribute to the processing of FlpHL–FRT DPCs, whereas the other DNA processing factors examined had limited effects under these conditions.

We also examined the effects of inhibiting additional DNA damage response and protein-processing pathways. Inhibition of ATM with AZD1390 did not increase FlpHL–FRT DPC accumulation (Fig. 2J). Similar results were obtained with inhibition of ATR with VE821 or DNA-PK with AZD7648, which produced little change in FLAG-FlpHL enrichment (Fig. 2K). Inhibition of the p97 unfoldase complex with CB-5083 resulted in a modest increase in DPC signal, although this effect was not statistically significant (Fig. 2L). Together, these results identify TDP1, MUS81, and XPF as factors whose loss increases FlpHL–FRT DPC accumulation, while perturbation of several other DNA repair and damage response pathways produced smaller or more variable effects under the conditions examined.

#### MRE11 depletion and inhibition increase FlpHL–FRT DPC accumulation

We next examined whether MRE11 contributes to the processing of FlpHL–FRT DPCs. MRE11 has known nuclease activity toward protein-blocked DNA ends but has primarily been implicated in the removal of 5′ covalent adducts ^5,7,8^. MRE11 acts on DNA ends through its endonuclease as well as 3′-to-5′ exonuclease activities at protein-blocked DNA double-strand breaks in a manner stimulated by phosphorylated CtIP ^4–9,25^.

MRE11 was depleted in HEK-293 Flp-In T-REx cells using shRNA, and reduced MRE11 protein levels were confirmed by Western blotting throughout the doxycycline induction period (Fig. 3A). In the presence of MG132, MRE11-depleted cells showed increased FLAG-FlpHL enrichment at the FRT locus relative to control cells, with the largest difference observed after 8 h of doxycycline treatment (Fig. 3B).

**Figure 3.**
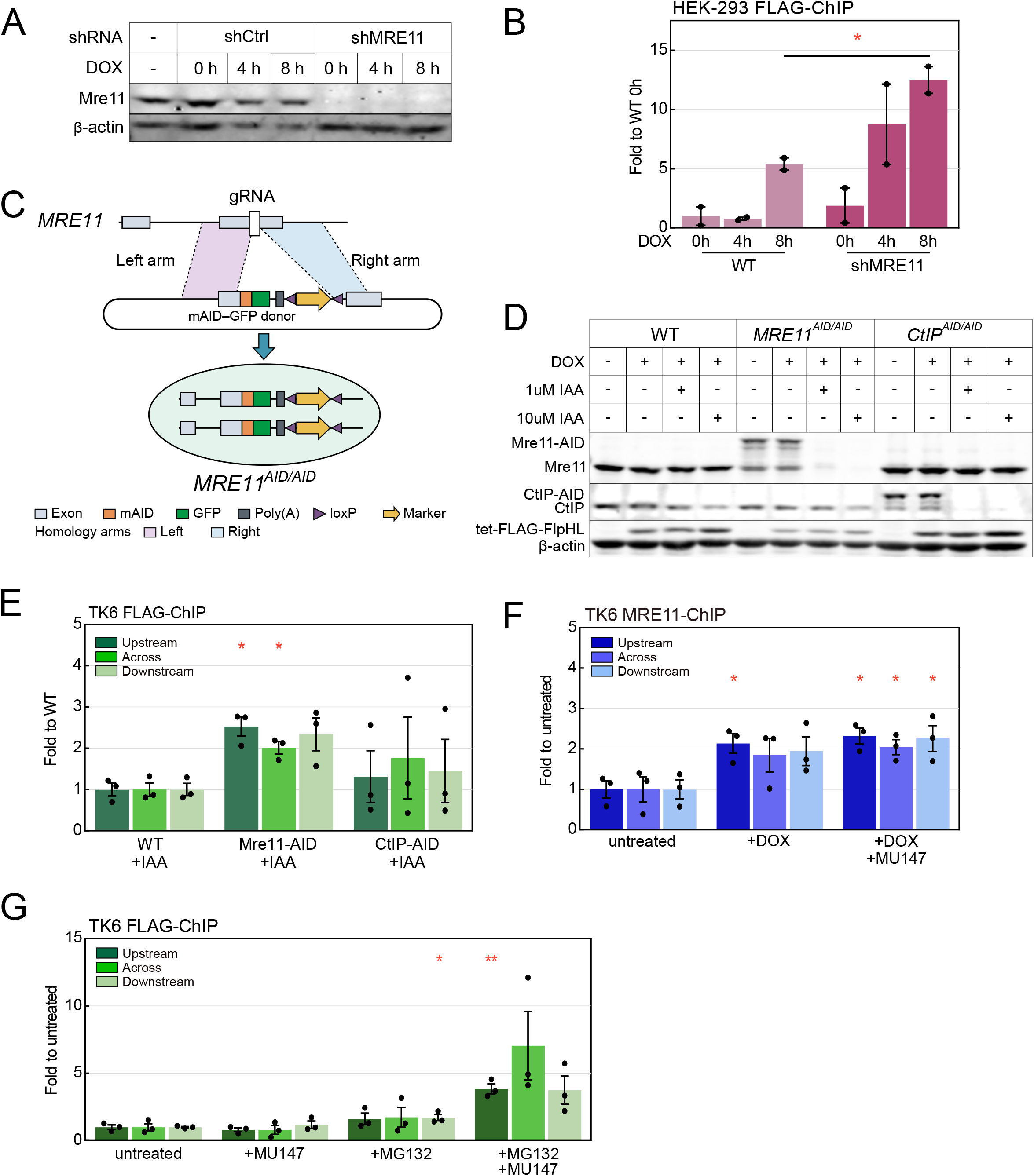
MRE11 depletion and inhibition increase FlpHL–FRT DPC accumulation. (A) Western blot analysis of MRE11 depletion in HEK-293 cells expressing control shRNA (shCtrl) or MRE11-targeting shRNA (shMRE11). Cells were treated with DOX (1 μg/mL) for the indicated times, and MRE11 protein levels were analyzed by immunoblotting. The leftmost lane represents parental cells without shRNA or DOX treatment. β-actin was used as a loading control. (B) ChIP-qPCR analysis of FlpHL–FRT DPC accumulation in control and MRE11-depleted HEK-293 cells shown in A. Cells were treated with DOX for the indicated times in the presence of MG132 (30 μM). FLAG-FlpHL associated with the integrated FRT site was measured by anti-FLAG ChIP-qPCR. Values were normalized to the 0-h shCtrl sample. Data represent two biological replicates; individual points represent biological replicates and bars show mean ± SEM. (C) Schematic diagram of *MRE11^AID/AID^* cell line construction. (D) Western blot validation of auxin-inducible depletion of MRE11 or CtIP in engineered TK6 cells carrying the FRT locus and doxycycline-inducible FLAG-FlpHL. WT, *MRE11^AID/AID^*, and *CtIP^AID/AID^* cells were pretreated with 5-Ph-IAA at the indicated concentrations for 18 h, followed by an additional 8 h in the continued presence of 5-Ph-IAA and DOX, where indicated. AID-tagged and endogenous MRE11 and CtIP species, together with FLAG-FlpHL, were analyzed by immunoblotting. β-actin was used as a loading control. (E) ChIP-qPCR analysis of FlpHL–FRT DPC accumulation in WT, MRE11-AID, and CtIP-AID TK6 cells following AID-mediated protein depletion. Cells were pretreated with 5-Ph-IAA (10 μM) for 18 h, followed by 8 h of DOX and MG132 treatment in the continued presence of 5-Ph-IAA. FLAG-FlpHL enrichment at the upstream, across, and downstream regions defined in Fig. 2A was measured by anti-FLAG ChIP-qPCR. Values for each primer set were normalized to the corresponding WT value. Data represent three biological replicates; individual points represent biological replicates and bars show mean ± SEM. (F) MRE11 ChIP-qPCR analysis at the engineered FRT locus following FlpHL induction and MRE11 inhibition. Cells were pretreated with or without MU147 (25 μM) for 18 h and then treated with DOX for 8 h in the continued presence or absence of MU147, as indicated. MG132 was present during the final 8 h in all conditions. Following formaldehyde crosslinking, MRE11-associated chromatin was immunoprecipitated, and enrichment at the upstream, across, and downstream regions defined in Fig. 2A was measured by qPCR. Values for each primer set were normalized to the corresponding untreated control. Data represent three biological replicates; individual points represent biological replicates and bars show mean ± SEM. (G) ChIP-qPCR analysis of FlpHL–FRT DPC accumulation in TK6 cells treated with the MRE11 inhibitor MU147 and/or MG132. All cells were treated with DOX for 8 h. Where indicated, cells were pretreated with MU147 (25 μM) for 18 h and maintained in MU147 during the subsequent 8 h treatment, while MG132 (30 μM) was added for the final 8 h. FLAG-FlpHL enrichment at the upstream, across, and downstream regions defined in Fig. 2A was measured by anti-FLAG ChIP-qPCR. Values were normalized to the corresponding untreated condition lacking MU147 and MG132. Data represent three biological replicates; individual points represent biological replicates and bars show mean ± SEM. For B and E–G, statistical significance was determined using a two-tailed Student’s *t* test assuming equal variance. Unless otherwise indicated by horizontal comparison lines, comparisons were made between each condition and the corresponding control for the same primer set. *, **, and *** indicate *P* < 0.05, *P* < 0.01, and *P* < 0.001, respectively; ns, not significant.

We also examined MRE11 function in TK6 cells by constructing an auxin-inducible degron cell line for MRE11 (Fig. 3C) and also using an equivalent line with an auxin-inducible degron CtIP ^26^. These lines have homozygous insertions of mAID into the endogenous MRE11 and CtIP loci, allowing for nearly complete removal of each protein with auxin. Here we used a codon-optimized version of OsTir1 with the engineered ligand 5-Ph-IAA ^27^. Treatment with 5-Ph-IAA reduced the corresponding levels of AID-tagged proteins, with near-complete depletion observed at 10 μM 5-Ph-IAA, while doxycycline induced FLAG-FlpHL expression in all three cell lines (Fig. 3D). We then measured FLAG-FlpHL enrichment following depletion of MRE11 or CtIP using the upstream, across, and downstream ChIP-qPCR primer sets described in Fig. 2. MRE11 depletion increased FLAG-FlpHL enrichment at all three regions relative to wild-type cells, with significant increases at the upstream and across regions (Fig. 3E). In contrast, CtIP depletion produced a more variable response and did not result in a significant increase in FLAG-FlpHL enrichment. These results are consistent with a role for MRE11 in FlpHL–FRT DPC processing, whereas the contribution of CtIP was not resolved under these conditions.

We next asked whether MRE11 becomes enriched at the FlpHL–FRT locus following DPC induction. Cells were treated with MG132 and doxycycline and subjected to formaldehyde crosslinking followed by MRE11 ChIP-qPCR. Doxycycline-induced FlpHL expression increased MRE11 association with the FRT locus relative to the untreated control (Fig. 3F), indicating that MRE11 is recruited to or retained at the FRT locus following FlpHL induction.

Finally, to examine the contribution of MRE11 nuclease activity to FlpHL–FRT DPC processing, we treated TK6 cells with the MRE11 nuclease inhibitor MU147 ^28^ in the presence or absence of MG132. Neither MU147 nor MG132 alone produced a substantial increase in FLAG-FlpHL enrichment in wild-type cells; however, combined treatment with MU147 and MG132 increased FLAG-FlpHL enrichment at all three regions, with the largest increase observed using the “across” primer set (Fig. 3G). This result is consistent with a contribution of MRE11 nuclease activity to FlpHL–FRT DPC processing under conditions in which proteasome activity is also inhibited.

### FlpHL processing in budding yeast

The levels of FlpHL DPC formation we observed in human cells are quantifiable but relatively low, likely because the system evolved to operate in fungal organisms. We turned to budding yeast to validate the role of MRE11 in FlpHL removal by inducibly expressing the protein in its native host, *S. cerevisiae* ^15^. The FRT target sequence was inserted upstream of the *ADE2* gene in a haploid W303 background, with FlpHL regulated by a galactose-inducible promoter (Fig. 4A). High levels of FLAG-tagged FlpHL were observed by 4 to 6 h following galactose addition, whereas no expression was seen in glucose (Fig. 4B).

**Figure 4.**
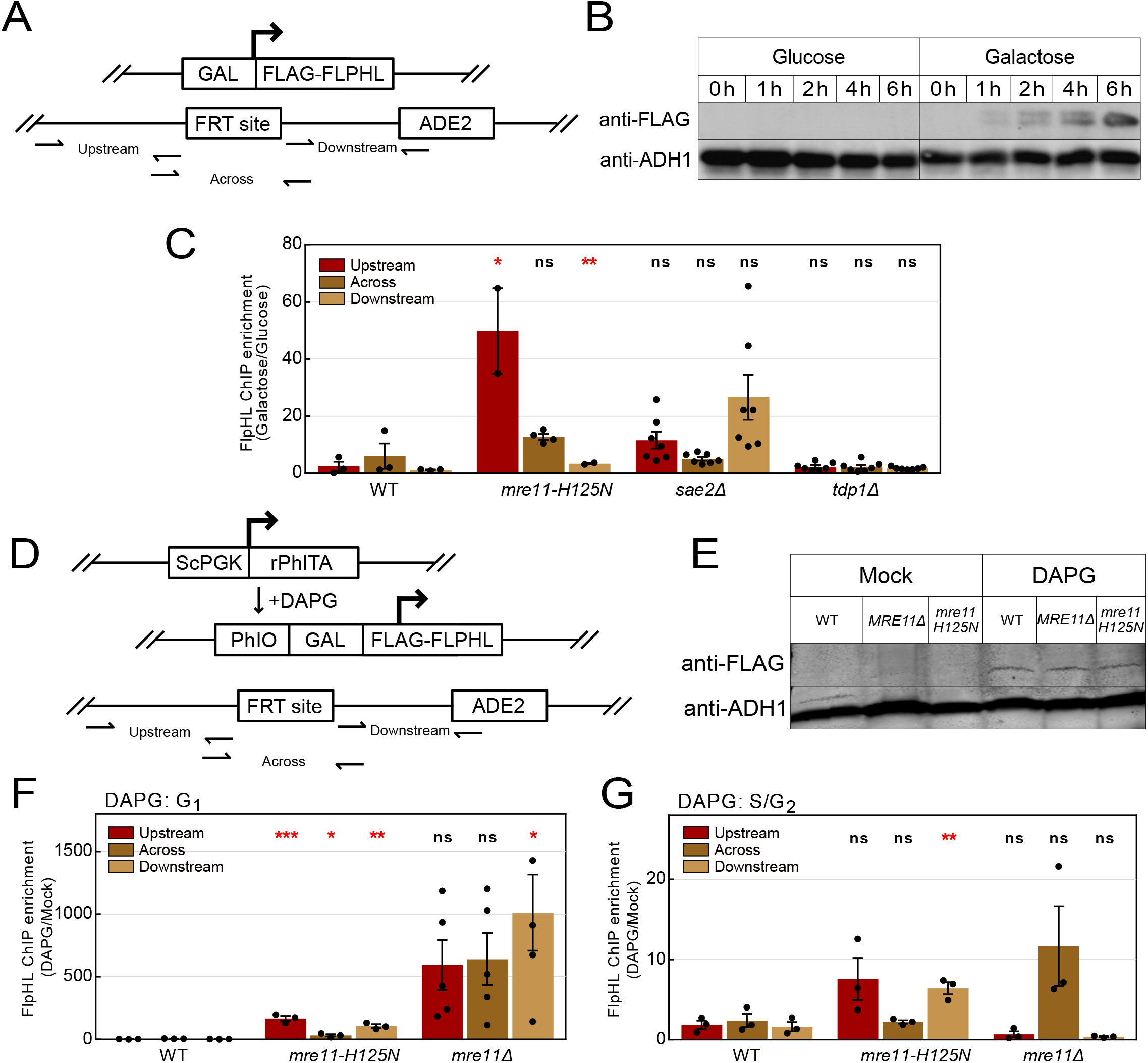
MRE11 nuclease activity promotes processing of Flp-H305L–FRT DPCs in *Saccharomyces cerevisiae*. (A) Schematic of the galactose-inducible Flp-H305L DPC system. Plasmid pTP4529 expresses 2×FLAG-tagged Flp-H305L under control of the *GAL1* promoter. Flp-H305L binds the FRT site inserted upstream of *ADE2* on the chromosome and forms a covalent DNA–protein crosslink at the FRT site. Small horizontal arrows indicate primer-binding sites defining the Upstream, Across, and Downstream amplicons used for ChIP-qPCR analyses. (B) Western blot showing induction kinetics of 2×FLAG-Flp-H305L. Cells were sampled at 0, 1, 2, 4, and 6 h after transfer to glucose- or galactose-containing medium. FLAG-Flp-H305L expression was detected by anti-FLAG immunoblotting, with Adh1 used as a loading control. (C) ChIP-qPCR analysis of Flp-H305L accumulation at the FRT locus in WT, *mre11-H125N*, *sae2Δ*, and *tdp1Δ* cells following glucose or galactose treatment for 6 h. Because Flp-H305L forms a stable covalent DNA–protein adduct through its catalytic reaction, no chemical crosslinking was performed before ChIP. FLAG-Flp-H305L enrichment at the Upstream, Across, and Downstream regions was measured by ChIP-qPCR. For each strain and primer set, enrichment in galactose-induced cells was calculated relative to the corresponding glucose-treated condition (Galactose/Glucose). Individual points represent biological replicates; bars show mean ± SEM. (D) Schematic of the DAPG-inducible Flp-H305L system. Plasmid pTP6004 expresses 2×FLAG-tagged Flp-H305L from a PhlF-responsive promoter. The PhlF-responsive transcriptional activator rPhlTA is expressed under the *S. cerevisiae PGK1* promoter, and addition of DAPG induces Flp-H305L expression. As in A, the FRT site is located upstream of *ADE2*, and the same Upstream, Across, and Downstream amplicons were used for ChIP-qPCR analyses. (E) Western blot showing DAPG-induced expression of 2×FLAG-Flp-H305L in WT, *mre11Δ*, and *mre11-H125N* cells. Cells were mock treated or treated with 50 µM DAPG for 2 h. FLAG-Flp-H305L expression was detected by anti-FLAG immunoblotting, with Adh1 used as a loading control. (F) ChIP-qPCR analysis of Flp-H305L accumulation at the FRT locus following nocodazole synchronization and release into DAPG-containing medium. WT, *mre11-H125N*, and *mre11Δ* cells were treated with nocodazole (15 µg/mL) for 2 h, followed by 50 µM DAPG for 30 min while cells remained nocodazole-arrested. Cells were then released from nocodazole into medium containing 50 µM DAPG and harvested 90 min after release. FLAG-Flp-H305L enrichment at the Upstream, Across, and Downstream regions was measured by ChIP-qPCR. For each strain and primer set, DAPG-induced enrichment was calculated relative to the corresponding uninduced condition. Individual points represent biological replicates; bars show mean ± SEM. (G) ChIP-qPCR analysis of Flp-H305L accumulation at the FRT locus following α-factor synchronization and release into DAPG-containing medium. WT, *mre11-H125N*, and *mre11Δ* cells were treated with α-factor (10 µg/mL) for 2 h, with arrest confirmed by shmoo formation. Cells were then treated with 50 µM DAPG together with an additional 5 µg/mL α-factor for 1 h while remaining arrested, released into medium containing 50 µM DAPG, and harvested 1 h after release. FLAG-Flp-H305L enrichment at the Upstream, Across, and Downstream regions was measured by ChIP-qPCR. For each strain and primer set, DAPG-induced enrichment was calculated relative to the corresponding uninduced condition. Individual points represent biological replicates; bars show mean ± SEM. For C, F, and G, statistical significance was determined using a two-tailed Student’s *t* test assuming equal variance. *, **, ***, and **** indicate *P* < 0.05, *P* < 0.01, *P* < 0.001, and *P* < 0.0001, respectively; ns, not significant. Exact *P* values are shown for selected comparisons that did not reach statistical significance.

ChIP-qPCR for FLAG-FlpHL was performed similarly to that in mammalian cells with no formaldehyde crosslinking, with variations in the procedure added for disruption of the yeast cell wall (Materials and Methods). For each strain and primer set, FlpHL enrichment following galactose induction was calculated relative to the corresponding glucose-treated condition. The qPCR results showed little accumulation in wild-type cells, while high levels of FLAG-FlpHL accumulation were observed following galactose induction in cells carrying the MRE11 nuclease-deficient H125N mutation, with the highest accumulation seen with the upstream primer set (Fig. 4C). Higher FlpHL accumulation was also observed in *sae2Δ* cells compared to wild-type, whereas *tdp1Δ* cells showed little difference between galactose and glucose conditions (Fig. 4C). These results suggest that MRE11 nuclease activity acts to process FlpHL–FRT DPCs in budding yeast and that Sae2 may also contribute to this function, although the FlpHL recovery in the *sae2Δ* line did not reach significance in our assays. The *mre11-H125N* phenotype is consistent with our results using an MRE11 nuclease inhibitor in human cells, whereas the lack of a detectable effect in tdp1Δ cells differs from the increased FlpHL accumulation observed in TDP1-deficient human cells.

To regulate FlpHL expression independently of growth conditions, we turned to an alternative system utilizing 2,4-diacetylphloroglucinol (DAPG) to induce a heterologous promoter via the DAPG-regulated transcriptional activator rPhlTA ^29^. rPhlTA was expressed via the yeast PGK1 promoter on a plasmid also containing FlpHL regulated by Phl operator sequences (Fig. 4D). Addition of DAPG to the culture induced FlpHL expression within 2 h in wild-type, *mre11-H125N*, and *mre11Δ* strains (Fig. 4E). We used this system to examine FlpHL accumulation in different cell-cycle phases.

Wild-type, *mre11-H125N*, and *mre11Δ* strains were synchronized with nocodazole, then treated with DAPG during release into G_1_ phase. For each strain and primer set, DAPG-induced FlpHL enrichment was calculated relative to the corresponding mock-treated condition. Under these conditions, high levels of FlpHL accumulation were observed in both *mre11-H125N* and *mre11Δ* strains compared with wild-type cells (Fig. 4F). Synchronization with α-factor was also performed in these strains, with DAPG added during and following release from α-factor into S/G_2_ phase. Recovery of FlpHL was overall lower under these conditions, but increased levels of DPC accumulation were observed in *mre11-H125N* and *mre11Δ* cells compared with wild-type cells (Fig. 4G).

### Mass spectrometry analysis identifies increased DNA-associated topoisomerases following MRE11 inhibition

The FlpHL system is useful for analysis of a site-specific 3ʹ DPC, but we also sought to extend this analysis to endogenous factors that form covalent linkages in human cells. To identify these factors, we developed a mass spectrometry workflow to isolate proteins covalently associated with genomic DNA from TK6 cells without chemical crosslinking. Cells were lysed under denaturing conditions with guanidine thiocyanate, followed by genomic DNA purification on a silica matrix and high-stringency washes. Proteins retained during DNA purification were digested on-column with trypsin, and the recovered peptides were analyzed by label-free mass spectrometry (Fig. 5A). We first examined TOP1 recovery in TDP1-deficient cells, as TDP1 is involved in the removal of TOP1–DNA covalent intermediates ^30,31^. TOP1 abundance was increased approximately fivefold in the DNA-associated fraction from TDP1-deficient cells compared with WT control samples (Fig. 5B). This result shows that this workflow can detect increased recovery of TOP1 covalent complexes (TOP1cc) under conditions associated with impaired TOP1–DNA adduct repair. Core histones were generally not recovered in this experiment (Supp. Table 2), indicating that tightly bound non-covalent associations are effectively removed by this strategy.

**Figure 5.**
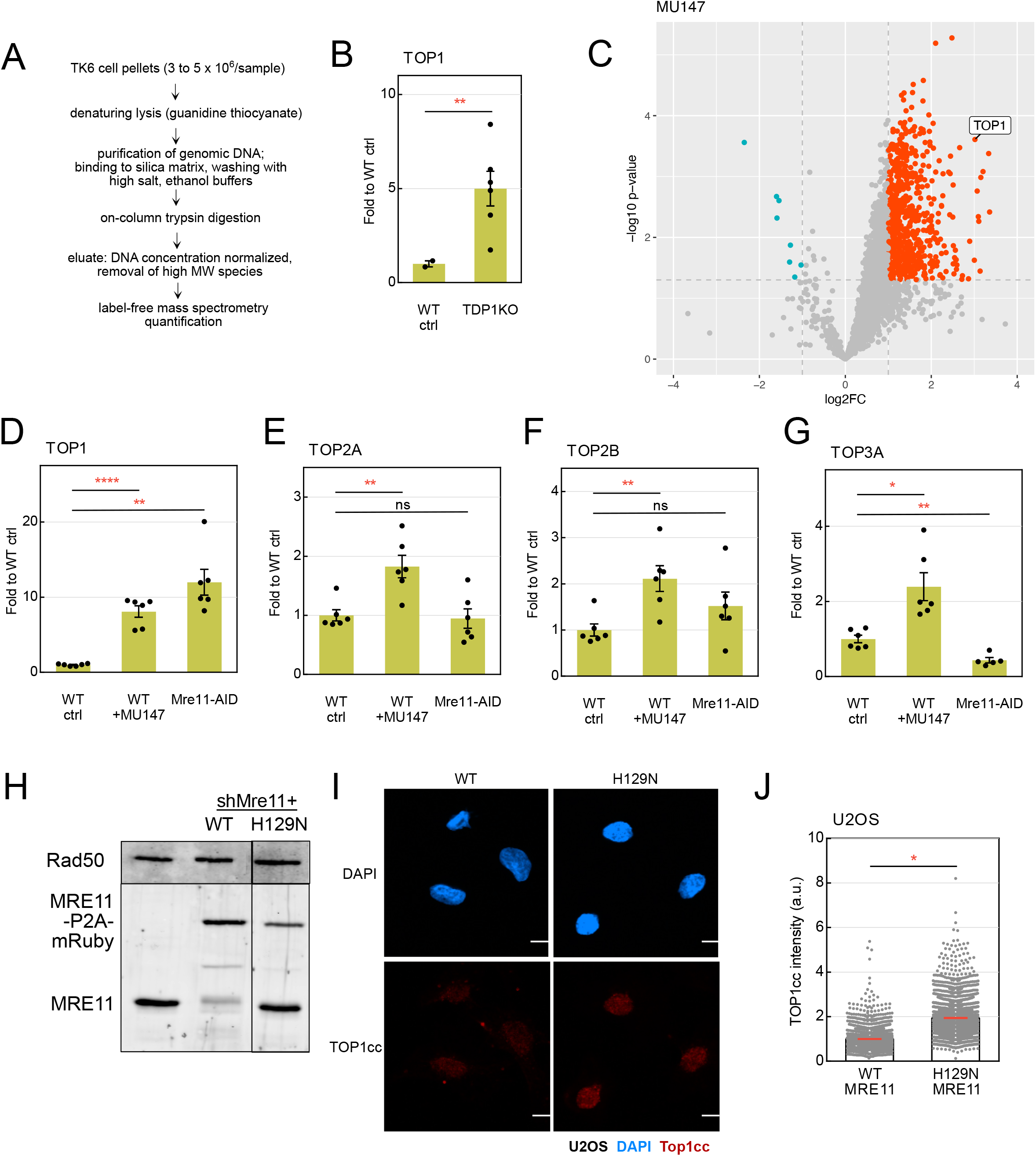
Mass spectrometry analysis identifies increased DNA-associated topoisomerases following MRE11 inhibition. (A) Schematic of the workflow used to isolate proteins covalently associated with genomic DNA for mass spectrometry. TK6 cell pellets were subjected to denaturing lysis with guanidine thiocyanate, followed by purification of genomic DNA using a silica matrix and sequential washes with high-salt and ethanol-containing buffers. Proteins retained during DNA purification were digested on-column with trypsin. Recovered peptides were processed for DNA concentration normalization and removal of high-molecular-weight species before label-free mass spectrometry analysis. No chemical crosslinking was performed. (B) TOP1 abundance in DNA-associated material isolated from WT and *TDP1−/−* TK6 cells treated with MG132 (30 μM). WT control cells lacked Flp-H305L expression and were treated with MG132 (30 μM), whereas TDP1−/− cells expressing Flp-H305L were treated with DOX (1 μg/mL) and MG132 (30 μM) for 8 h. TOP1 abundance was normalized to sample input and expressed relative to the WT control. Individual points represent biological replicates; bars show mean ± SEM. (C) Volcano plot comparing protein abundance in DNA-associated material isolated from WT TK6 cells treated with MU147 (25 μM) for 4 d versus untreated WT controls. Protein abundance was normalized to sample input, and differential abundance is plotted as log2 fold change (MU147 samples/untreated samples) versus −log10 p value. TOP1 is indicated. Positive log2 fold-change values represent proteins enriched following MU147 treatment. See Supp. Table 2. (D–G) Abundance of TOP1 (D), TOP2A (E), TOP2B (F), and TOP3A (G) in DNA-associated material from mass spectrometry analysis as in C. Samples were derived from untreated WT TK6 cells, WT TK6 cells treated with MU147 (25 μM) for 4 d, and MRE11-AID TK6 cells treated with 5-Ph-IAA (10 μM) for 5 d to deplete MRE11. Protein abundance was normalized to sample input and expressed relative to the WT control. Individual points represent biological replicates; bars show mean ± SEM. (H) Western blot analysis of MRE11 in vector-control, WT MRE11-P2A-mRuby, and MRE11 H129N-P2A-mRuby U2OS cells following doxycycline treatment. The upper band corresponds to MRE11-P2A-mRuby, whereas the lower band corresponds to MRE11 with P2A-mediated ribsome skipping. RAD50 protein levels were blotted and analyzed in parallel as a control. (I) Representative immunofluorescence images of TOP1 covalent complexes (TOP1cc) in asynchronous U2OS cells expressing shMRE11-resistant MRE11-WT or MRE11-H129N. Cells were treated with doxycycline for 48 h to induce expression of the indicated MRE11 construct while endogenous MRE11 was depleted by shRNA. Cells were subsequently fixed and stained for TOP1cc. DAPI was used to visualize nuclei. (J) Quantification of TOP1cc fluorescence intensity in MRE11-WT- and MRE11-H129N-expressing U2OS cells. Each gray point represents an individual cell, and TOP1cc intensity is expressed in arbitrary units (a.u.). Data shown were obtained from three independent experiments per condition. For B and D–G, statistical significance was determined using two-tailed Welch’s *t* tests assuming unequal variance. For J, statistical significance was determined using the average values from 3 independent experiments, the means were analyzed using a two-tailed Student’s *t* test assuming equal variance. *, **, ***, and **** indicate *P* < 0.05, *P* < 0.01, *P* < 0.001, and *P* < 0.0001, respectively; ns, not significant. More than 500 cells were analyzed in each replicate.

We next asked whether MRE11 inhibition or depletion alters the association of endogenous factors with genomic DNA using TK6 cells without FlpHL expression. Proteome-wide analysis of WT TK6 cells treated with the MRE11 inhibitor MU147 identified TOP1 among the proteins enriched in the DNA-associated fraction relative to untreated WT cells (Fig. 5C). Quantification of TOP1 abundance showed an approximately eightfold increase following MU147 treatment and an approximately 12-fold increase following MRE11 depletion in MRE11-AID cells compared to wild-type untreated cells (Fig. 5D). These results indicate that both MRE11 inhibition and depletion are associated with increased recovery of endogenous DNA-associated TOP1 in the absence of exogenous DNA damage.

We also examined whether other topoisomerases showed similar changes in this dataset. MU147 treatment increased the abundance of TOP2A, TOP2B, and TOP3A in the DNA-associated fraction by approximately twofold (Fig. 5E–G). In contrast, MRE11 depletion did not produce a corresponding increase in TOP2A or TOP2B, while TOP3A abundance was reduced relative to WT cells. Thus, although MRE11 inhibition increased the recovery of all four topoisomerases examined, MRE11 depletion was associated with increased TOP1 recovery but not with a similar increase in the other topoisomerases.

To quantitate endogenous TOP1 covalent complexes using an orthogonal method, we examined TOP1cc levels by immunofluorescence in U2OS cells expressing shMRE11-resistant wild-type MRE11-P2A-mRuby or the nuclease-deficient H129N MRE11-P2A-mRuby while endogenous MRE11 was depleted. Western blot analysis confirmed expression of the wild-type and mutant MRE11 proteins following doxycycline induction, although the P2A-mediated ribosome skipping was variably efficient here, yielding both MRE11-P2A-mRuby and untagged MRE11 forms of the protein (Fig. 5H). TOP1cc levels were increased in cells expressing MRE11-H129N compared with cells expressing MRE11-WT (Fig. 5I, J). These results provide independent evidence that impaired MRE11 nuclease activity is associated with increased accumulation of endogenous TOP1cc. Together, these results show that MRE11 inhibition and depletion have distinct effects on the abundance of DNA-associated topoisomerases, with TOP1 increasing under both conditions, and that MRE11 nuclease deficiency is associated with increased endogenous TOP1cc accumulation.

## Discussion

We established an inducible Flp-H305L–FRT system to examine the processing of a defined 3′-phosphotyrosyl DNA–protein crosslink in mammalian cells and budding yeast. Our results identify MRE11 nuclease activity as an important determinant of Flp-DPC persistence, with supporting genetic evidence from investigation of MRX in budding yeast. Multiple additional repair factors, including phosphotyrosyl-DNA phosphodiesterases and structure-specific nucleases, also contribute to Flp-DPC resolution in combination with proteasomal activity, suggesting that this lesion can be processed through parallel pathways. Finally, the accumulation of endogenous topoisomerase-associated covalent cleavage complexes following MRE11 perturbation indicates that the role of MRE11 extends to 3′ TOP1-DNA adducts in addition to its well-known role in processing Spo11 and other 5’ DPCs.

### FlpH305L as a site-specific model for 3′-phosphotyrosyl DPC processing

FlpH305L generates a stable, defined 3′-phosphotyrosyl protein–DNA adduct by trapping the normally transient covalent intermediate of the Flp recombination reaction ^12,13^. This chemistry makes the Flp-nick system useful for studying how a site-specific protein-bound nick affects replication, recombination, and DPC clearance, particularly in yeast, and more recently in mammalian cells ^14–19^. In our study, we exploited the Flp lesion as a direct readout of DPC persistence rather than as a trigger for downstream repair outcomes, using FLAG-FlpHL enrichment at the FRT locus to monitor the covalent adduct in mammalian cells and in budding yeast. This distinction allowed us to compare the fate of the covalent adduct itself across the two cellular systems and to test whether conserved DPC-processing factors act similarly on the same chemically defined lesion. Monitoring the DPC directly is an important consideration here, as the MRN(X) complex is involved in homologous recombination that occurs subsequent to DPC removal as well, so measurement of HR outcomes following DPC formation does not necessarily show a role in DPC processing.

Quantitation of the Flp-HL DPC lesion in mammalian cells revealed increases in Flp-DPC accumulation in MRE11-deficient cells, as well as cells lacking TDP1, MUS81, and XPF. These increases in Flp accumulation were generally modest but reproducible, and were only observed in the absence of proteasome function (Fig. 2D, H; Fig. 3B, E, G). In budding yeast, by contrast, genetic disruption of the MRX–Sae2 pathway produced substantially larger changes in FlpHL accumulation, which was observed without proteasome disruption. Previous Flp-nick studies have shown that the same lesion can be processed differently depending on replication, proteolysis, and repair-pathway context ^15–18^, and our results extend this principle across species. We speculate that the stronger dynamic range in yeast is likely due to the more efficient formation of FlpHL DPCs in its native host.

### MRE11 nuclease activity contributes to 3′-protein adduct processing

Our data add several lines of evidence that MRE11 contributes to the processing of 3′-protein adducts themselves, rather than acting only after conversion of the lesion into a conventional DNA break. First, MRE11 depletion increased FlpHL retention at the FRT locus in both HEK-293 and TK6 cells (Fig. 3B, E), and MRE11 was itself enriched at the lesion following FlpHL induction and accumulated further when its nuclease activity was inhibited (Fig. 3F). Catalytic inhibition of MRE11 also increased FlpHL persistence, particularly when proteasome activity was simultaneously blocked (Fig. 3G), arguing that MRE11 nuclease activity contributes to lesion processing in parallel with or after proteolytic remodeling. These observations extend previous biochemical studies of protein-blocked DNA ends, which largely defined MRE11 nuclease activity on 5′-linked or terminal protein adducts ^4,5,8^, to a genetically defined 3′-phosphotyrosyl DPC in cells. Importantly, the requirement for MRE11 was conserved in budding yeast, where the nuclease-defective mre11-H125N allele caused strong FlpHL accumulation and loss of Sae2 similarly impaired lesion processing (Fig. 4C). This is consistent with the established role of Sae2 in stimulating MRX endonuclease activity at protein-blocked DNA ends ^4,6^ but contrasts with the weaker and more variable effect of CtIP depletion in our mammalian system (Fig. 3E) and a less obvious role for CtIP in vitro on streptavidin-DNA adducts ^5^. Together, these results suggest that the catalytic activity of MRE11 is a conserved determinant of Flp-DPC removal, whereas the requirement for its accessory factors may depend on substrate architecture or cellular context.

Working with MRE11 is challenging because it is essential in dividing mammalian cells ^32^, and depletion of the vast majority of the protein is necessary for loss of nuclease function. Reduction of MRE11 to the point where it is not visible by western blotting is not necessarily sufficient for elimination of its function, as we have observed that even 5% of normal MRE11 levels is functionally equivalent to wild-type cells in some assays. The AID-MRE11 alleles used here in TK6 cells allow for nearly complete and inducible loss of the protein, and the inhibitor MU147 also provides a means to block the catalytic activity of MRN in a cellular context.

### Protein remodeling and lesion architecture may shape MRE11-dependent DPC processing

In this work we observed that proteasome inhibition increased the steady-state accumulation of FlpHL at the FRT locus in mammalian cells (Fig. 1C, D), and inhibition of MRE11 nuclease activity produced a substantially stronger increase in Flp-DPC persistence when combined with MG132 than with MU147 alone (Fig. 3G). Inhibition of the AAA+ ATPase p97, an ATP-driven protein unfoldase ^33^, also produced a modest increase in FlpHL accumulation (Fig. 2L). These observations are consistent with a model in which protein remodeling and nuclease-dependent processing constitute parallel and at least partially redundant routes for DPC resolution. Proteasome-mediated processing of ubiquitin-modified DPC substrates has been demonstrated in Xenopus extracts and in mammalian cells ^34–38^.

Recent work characterizing the removal of a covalent OGG1 DPC in human cells showed proteasome-dependent nucleotide excision repair in parallel with a homologous recombination-mediated pathway that is proteasome-independent ^35^. MRN function was not explicitly tested in this study but the results are consistent with our observations of a proteasome-independent, MRN-dependent pathway of DPC removal. Interestingly, ubiquitination was essential for the homologous recombination pathway of DPC removal in the OGG1 DPC assay even though proteasome function was not. Multiple groups have also shown replication-dependent functions of the SPRTN metalloprotease that are promoted by ubiquitin and target various types of DPCs including TOP1 ^36,38^.

MRE11 acts as an endonuclease in a manner that requires a protein block ^5^, thus it is likely that recognition of the protein adduct itself as a lesion promotes MRE11 endonuclease activity on the adjacent, covalently attached DNA. This mechanism would theoretically occur independently of protein degradation, in contrast to TDP1 which requires proteasome-mediated removal of most of a lesion prior to the direct reversal of protein-DNA crosslinks^3^. An important unresolved question is the DNA structure on which MRE11 acts during Flp-DPC processing. Although FlpH305L initially generates a protein-linked nick, replication or transcription through the lesion could generate secondary DNA structures that differ substantially from the original adduct. Previous Flp-nick studies have demonstrated replication-dependent conversion of the lesion into broken-fork intermediates that include the Flp 3’ DPC on single-strand overhangs and double-strand breaks ^14,19^. We know from biochemical studies that MRE11 activity toward protein-blocked double-strand breaks is strongly influenced by DNA-end architecture and can act on 3’ protein lesions ^5,9^. Based on this in vitro evidence we hypothesize that replication-induced remodeling of Flp 3’ DPCs into DPC-associated double-strand breaks likely generates the optimal substrate for MRN(X) processing. Our observation of FlpHL DPCs during S/G_2_ phase in budding yeast is consistent with this hypothesis; however, we also found very high levels of DPC formation in G_1_ phase. We do not know the architecture of Flp DPCs in G_1_ and how MRX targets this lesion in this scenario. It is possible in yeast that Flp DPC formation occurs efficiently on both strands, thus generating double-strand breaks independent of replication. The recovery of Flp-bound DNA fragments in this cell cycle phase is highest using the "upstream" and "downstream" primer pairs compared to the "across" primer set, likely an indication that there are breaks on both strands. It is not clear why the recovery of stable FlpHL DPCs appears to be much higher in MRE11 mutant cells in G_1_ compared to G_2_ phase; we hypothesize that S phase-specific redundant pathways such as WSS1 (Spartan) may act redundantly during S phase to remove the lesion.

### MRE11 perturbation alters the association of multiple topoisomerases with DNA

Our mass spectrometry analysis of tightly bound proteins on genomic DNA as well as immunofluorescence-based quantitation of TOP1cc suggests that MRE11 plays a direct role in removal of endogenous 3’ TOP1 lesions in mammalian cells. This connection is particularly important because prior work on 3′ bulky adducts has suggested that MRN can promote resection without an absolute requirement for MRE11 nuclease activity, whereas other studies have implicated MRE11 directly in TOP1-associated damage ^6,9,10^. Our results therefore support a model in which MRE11 can contribute directly to the processing of 3′-protein adducts, but they do not yet define whether MRE11 incises DNA adjacent to the intact Flp-DPC, acts after partial proteolysis, or encounters a secondary DNA structure generated during replication or transcription.

In addition to the strong increase in DNA-associated TOP1, MRE11 inhibition increased recovery of TOP2A, TOP2B, and TOP3A by approximately twofold in human TK6 cells (Fig. 5D–G). These enzymes generate structurally distinct catalytic intermediates: TOP1 forms a 3′-phosphotyrosyl single-strand cleavage complex, whereas TOP2A and TOP2B form 5′-phosphotyrosyl double-strand cleavage complexes, and TOP3A is a type IA topoisomerase that forms a 5′-phosphotyrosyl single-strand intermediate on single-stranded or recombination-associated DNA structures ^11,39,40^. The enrichment of multiple topoisomerases following MRE11 inhibition is therefore consistent with MRE11 contributing to the processing of a broader range of protein-linked DNA structures. This interpretation is supported by previous work demonstrating MRE11-dependent removal of 5′-linked TOP2 cleavage complexes and, more recently, TOP3A cleavage complexes, for which MRE11 functions in parallel with a SPRTN–TDP2 pathway ^7,8,40^.

MRE11 inhibition and depletion produced distinct effects on DNA-associated topoisomerases (Fig. 5D–G). In our AID system, MRE11 protein was strongly depleted (Fig. 3D), whereas MU147 inhibits MRE11 endonuclease and exonuclease activities while leaving the protein and MRN complex present ^28^. Thus, MU147-treated cells may retain noncatalytic functions of MRE11, including structural or DNA-associated roles, that are lost upon MRE11 depletion. This difference could contribute to the divergent TOP2A, TOP2B, and TOP3A responses observed between the two conditions. The consistent increase in TOP1 under both perturbations (Fig. 5D), together with increased TOP1 cleavage complexes in MRE11 nuclease-deficient cells (Fig. 5I, J), provides the strongest evidence for a role of MRE11 in endogenous TOP1 lesion processing.

In summary, we show that MRE11 contributes to the processing of a defined 3′-phosphotyrosyl DNA–protein crosslink in both mammalian cells and budding yeast. Depletion or inhibition of MRE11 increases persistence of the FlpH305L adduct, and the requirement for MRE11 nuclease activity, together with recruitment of MRE11 to the lesion and the corresponding phenotype of nuclease-defective MRE11 in yeast, supports a direct role for MRE11 in DPC processing. Our genetic analyses further indicate that MRE11 functions within a broader network that includes phosphotyrosyl-DNA phosphodiesterases, structure-specific nucleases, and protein-remodeling pathways, while analysis of endogenous topoisomerases suggests that MRE11 activity may extend to multiple classes of covalent topoisomerase intermediates. The precise DNA structure recognized by MRE11 and the relationship between nuclease-dependent processing and proteolytic remodeling remain unresolved. Defining these intermediates, and determining how pathway choice is influenced by lesion architecture and cellular context, will be important directions for future studies.

## Acknowledgments

We acknowledge CPRIT grant 13023012 to T.P. and helpful advice from colleagues.

## Author Contributions

Conceptualization: Y.Y., T.P., Data curation: Y.Y., Formal analysis: Y.Y., N.E., Funding acquisition: T.P., Investigation: Y.Y., S.K., B.Y., N.E., H.B., O.E., Methodology: Y.Y., S.K., B.Y., N.E., H.B., O.E., Project Administration: T.P., Resources: L.S., H.S., J.X., S.T., Y.Y., T.P., Supervision: T.P., Validation: Y.Y., S.K., B.Y., N.E., H.B., O.E., Visualization: Y.Y., S.K., B.Y., N.E., H.B., Writing original draft: Y.Y., T.P., Writing review and editing: Y.Y., T.P., S.K., B.Y., N.E.

## Supplemental Information

Supplementary Table 1: list of cell lines, yeast strains, plasmids, and primers. Supplementary Table 2: mass spectrometry data

## Materials and Methods

### Cell culture

Human cell lines used in this study included HEK-293T, U2OS T-REx Flp-In, HEK-293 Flp-In T-REx, and TK6 cells (Supp. Table 1). HEK-293T, U2OS, and HEK-293 cells were maintained in DMEM supplemented with 10% tetracycline-negative fetal bovine serum and 1% penicillin-streptomycin. TK6 cells were maintained in RPMI-1640 supplemented with 10% tetracycline-negative fetal bovine serum, 1% penicillin-streptomycin, and 1 mM sodium pyruvate. Cells were cultured at 37°C in a humidified incubator containing 5% CO₂ unless otherwise indicated.

Construction of PARP-deficient TK6 cells: The *PARP1−/−* TK6 line was generated by Dr. Liton Saha by targeting exon 3 of PARP1 using CRISPR/Cas9 together with targeting constructs carrying Neo and Puro selection markers. Candidate clones were screened by PCR, and loss of PARP1 protein expression was confirmed by Western blotting. The *PARP1−/− PARP2+/−* line was subsequently generated by introducing PARP2-targeting constructs into *PARP1−/−* TK6 cells. Exon 3 of PARP2 was targeted using a guide RNA directed to this region together with targeting constructs carrying Hygro and HisD selection markers, and candidate clones were screened by PCR.

Construction of Auxin-inducible degron TK6 cells: For MRE11-AID/AID TK6 cell generation, left and right arms with MRE11 genomic sequences were cloned separately into pBS-mAID-GFP_NEO (pMK286, a gift from Masato Kanemaki; Addgene #72824) and an Mre11 gRNA sequence (5ʹ-CTGAGAAACATGCAAGATACAGG-3ʹ) was cloned into pX330-gRNA (a gift from Charles P. Lai; Addgene #158973). Each MRE11 donor plasmid contained an mAID–GFP cassette flanked by left and right *MRE11* homology arms of approximately 1 kb each. The two donors carried loxP-flanked selectable cassettes conferring neomycin resistance (Neo^R^) or hygromycin resistance (Hygro^R^), respectively^41^. The primers used to amplify the homology arms included MRE11-LA-F, MRE11-LA-R, MRE11-RA-F, and MRE11-RA-R (see Supp. Table 1). The 3 plasmids were transfected into TK6 cells with NEON (Thermo Fisher, 1350V, 10 ms, 3 pulses). After a two-day incubation, cells were seeded in 96-well plates with antibiotics (Neo and Hygro) to isolate single clones. The genotype was confirmed by PCR and Western blotting. The CtIP-AID/AID cells were generated as previously described ^26^. In both cell lines, pTP5741 was first introduced to express codon-optimized OsTir1 linked to P2A-eGFP. GFP-positive cells were isolated by fluorescence-activated cell sorting (Sony MA900), with non-transduced cells used as gating controls. Cells were then sequentially transduced and selected with lentiviruses pTP5622 (FRT, blasticidin resistance) and either pTP5620 (FlpHL, puro resistance) or pTP6101 (codon-optimized FlpHL, puro resistance). No significant functional differences were observed comparing pTP5620 and pTP6101.

### Lentiviral packaging and production

HEK-293T cells were used for lentivirus production using a three-plasmid packaging system consisting of the lentiviral expression vector, pCMB-VSV-G envelope plasmid (gift from Bob Weinberg, Addgene plasmid #8454) and Δ8.9 packaging plasmid ^42^. Transfections were performed using FuGENE HD Transfection Reagent (Promega, E2312) in Opti-MEM (Gibco). HEK-293T cells were seeded in 6-well plates and transfected at >90% confluence. For each well, 100 ng VSV-G plasmid, 500 ng Δ8.9 packaging plasmid, 1.2 µg lentiviral vector, and 7 µL FuGENE HD were combined in 100 µL Opti-MEM and incubated for 30–45 min at room temperature before addition to the cells. Lentiviral supernatants were collected at 48 and 72 h after transfection, pooled, filtered through a 0.45-µm syringe filter, aliquoted, and stored at −80°C.

### Lentiviral transduction of adherent cells

For U2OS and HEK-293 Flp-In cells, cells were seeded in 6-well plates and transduced at approximately 50% confluence by replacing 1 mL of the 2-mL culture volume with 1 mL lentiviral supernatant. After overnight incubation, viral medium was replaced with fresh complete medium. Antibiotic selection was initiated approximately 48 h later using 200 µg/mL hygromycin. Non-transduced cells were maintained in parallel as selection controls. The pTP4883 construct was used to generate tetracycline-inducible FLAG/BirA-Flp-H305L-expressing U2OS and HEK-293 Flp-In cells.

### Lentiviral transduction and generation of Flp-inducible TK6 cells

For TK6 transduction, cells were adjusted to 1 × 10⁶ cells/mL. One milliliter of cell suspension was combined with 0.5 mL fresh medium and 1 mL lentiviral supernatant in a 6-well plate. Plates were sealed with breathable film and centrifuged at 800 × g for 1.5 h at room temperature. Following overnight incubation, cells were washed with PBS, resuspended in fresh medium, and transferred to T25 flasks. Antibiotic selection was initiated 48 h after transduction. Selection agents included 10 µg/mL blasticidin, puromycin at genotype-optimized concentrations, and 200 µg/mL hygromycin as appropriate.

To generate Flp-inducible TK6 lines, cells were first transduced with pTP5622, which introduces a 34-bp FRT site within a heterologous transgene, and selected with blasticidin. Cells were subsequently transduced with a tetracycline-inducible Flp-H305L construct (pTP5620) and selected with puromycin. A codon-optimized derivative (pTP6101) was used in the experiments shown in Fig. 2D, Fig. 2K, and Fig. 3F.

### MRE11 shRNA depletion and complementation

For MRE11 depletion experiments in HEK-293 cells, pTP4112 was used to express MRE11-targeting shRNA from an eGFP-2A-Puro lentiviral backbone. The matched control vector pTP4114 was derived from pTP4112 by removal of the MRE11 shRNA cassette. In U2OS cells, wild-type and H129N Mre11-P2A-mRuby lentivirus constructs (pTP6166 and pTP6064, respectively) were used to express MRE11. These constructs also contain an Mre11-targeted shRNA driven by the H1 promoter ^43^ (see Supp. Table 1).

### *Saccharomyces cerevisiae* strains and plasmids

All experiments were performed using W303a *S. cerevisiae* strains (see Supp. Table 1 for details).

### Primers

See Supp. Table 1.

### Mammalian cell treatments

Flp-H305L-mediated DNA–protein crosslink formation was induced by addition of doxycycline hydrochloride (Fisher BioReagents, BP2653-1) to the culture medium at a final concentration of 1 µg/mL for the indicated times. Where indicated, the proteasome inhibitor MG132 (Cayman Chemical, 10012628) was added at a final concentration of 30 µM concurrently with doxycycline and maintained throughout the indicated treatment period. For pharmacological inhibition of MRE11 nuclease activity, MU147, provided by the Nikulenkov laboratory (Nikulenkov et al., 2025) or obtained commercially (MedChemExpress HY-170844), was used at 25 µM. Cells were pretreated with MU147 for approximately 18 h, and the compound was maintained during the subsequent indicated doxycycline treatment, with MG132 where indicated.

For inhibition of DNA damage response pathways, AZD1390 (Selleck Chemicals, S8680) was used at 1 µM and veliparib (Selleck Chemicals, S1004) at 10 µM. Cells were pretreated with either compound for approximately 18 h, and the inhibitors were maintained during the subsequent 8-h doxycycline/MG132 treatment. VE-821 (Tocris Bioscience, F8130) was used at 1 µM, AZD7648 (MedChemExpress, HY-111783) at 1 µM, and CB-5083 (ApexBio, B6032) at 2.5 µM during the 8-h doxycycline/MG132 treatment.

For auxin-inducible degron experiments, 5-Ph-IAA (MedChemExpress, HY-134653) was added at a final concentration of 10 µM approximately 18 h before DPC induction and maintained throughout the subsequent doxycycline/MG132 treatment. Where indicated, lower concentrations of 5-Ph-IAA were used to assess depletion efficiency, as specified in the corresponding figure. Unless otherwise indicated, mammalian-cell treatments were performed at 37°C with 5% CO₂. Cells were subsequently harvested for ChIP-qPCR, immunoblotting, or other analyses as described below.

### yeast induction and synchronization

For experiments using the GAL1-regulated Flp-H305L construct, cells were grown overnight in −LEU medium containing 2% glucose, washed twice with sterile water, and transferred to −LEU medium containing 2% raffinose for overnight growth at 30°C. Cultures were diluted the following day to an OD₆₀₀ of 0.1 in 200 mL −LEU medium containing 2% raffinose and grown to an OD₆₀₀ of approximately 0.4–0.5. Cultures were then divided into glucose- and galactose-treated groups. Galactose-dependent Flp-H305L induction and the subsequent glucose chase were performed for the indicated durations. For strains carrying the DAPG-inducible Flp-H305L system, 2,4-diacetylphloroglucinol (DAPG; Cayman Chemical, 16345) was added at a final concentration of 50 µM for the indicated times. Parallel mock-treated cultures received no DAPG. For nocodazole synchronization experiments, cells were treated with nocodazole (Sigma-Aldrich, M1404) at 15 µg/mL for 2 h. DAPG was then added at 50 µM for 30 min while cells remained nocodazole-arrested. Cells were subsequently released from nocodazole into medium containing 50 µM DAPG and harvested 90 min after release.

For α-factor synchronization experiments, cells were treated with α-factor mating pheromone (GenScript, RP01002) at 10 µg/mL for 2 h, with arrest confirmed visually by shmoo formation. DAPG was then added at 50 µM with an additional 5 µg/mL α-factor for 1 h. Cells were released from α-factor arrest into medium containing 50 µM DAPG and harvested 1 h after release.

### Western blot analysis: mammalian cells

Western blotting was performed to assess Flp-H305L expression and to validate depletion of MRE11 or CtIP. Approximately 1 × 10⁶ PBS-washed cells were harvested following the indicated treatments and resuspended directly in 200 µL of 1× SDS sample buffer. Samples were heated at 100°C for 10 min and stored at −20°C until analysis. Proteins were resolved on 8%, 10%, or 12% SDS-polyacrylamide gels, depending on protein molecular weight, and transferred to PVDF membranes using either semi-dry or wet transfer. PVDF membranes were pre-wetted in methanol and equilibrated in transfer buffer before use. Wet transfers were performed at 180 V and 0.4 A for 3 h at 4°C or overnight at 0.2 A. Membranes were washed in PBST (PBS containing 0.1% Tween-20) and blocked in LI-COR blocking buffer for 1 h at room temperature. Primary antibodies were incubated overnight at 4°C with gentle agitation. The following primary antibodies were used: anti-DDDDK-tag monoclonal antibody (MBL, clone M185-3L) for detection of FLAG-tagged Flp-H305L; anti-β-actin monoclonal antibody (ABclonal, C4, AC026) as a loading control; rabbit polyclonal anti-MRE11 antibody (Thermo Fisher Scientific, PA3-16527); and anti-CtIP monoclonal antibody (Active Motif, clone 14-1, 61141).

Following primary antibody incubation, membranes were washed with PBST and incubated for 1 h at room temperature in the dark with fluorescent secondary antibodies: Alexa Fluor 680 anti-rabbit IgG (Invitrogen, A21109) or Alexa Fluor Plus 800 goat anti-mouse IgG (H+L) (Invitrogen, A32730), as appropriate. Blots were imaged using a LI-COR Odyssey CLx system. Signal intensities were quantified using LI-COR Image Studio software or ImageJ where indicated.

### Yeast expression and lysis

For Flp-H305L expression time-course analysis and DAPG-induction analysis, 25 OD600 units of cells were harvested at the indicated time points before and after induction. Cells were lysed in 200 µL of the same lysis buffer used for ChIP, supplemented with protease inhibitors, by glass-bead disruption. Protein concentration was determined using Bradford reagent, and 50 µg of total protein was resolved per lane on 12% SDS-polyacrylamide gels. Proteins were transferred to PVDF membranes by wet transfer using Tris-glycine transfer buffer containing 10% methanol. 2×FLAG-tagged Flp-H305L was detected using mouse monoclonal anti-FLAG M2 antibody (Sigma-Aldrich, F1804) at a 1:2,000 dilution, and Adh1 was detected as a loading control using rabbit anti-Adh1 antibody (Abcam, ab20994) at a 1:5,000 dilution. Membranes were subsequently incubated with Alexa Fluor Plus 800 goat anti-mouse IgG (H+L) secondary antibody (Thermo Fisher Scientific, A32730) or Alexa Fluor 680 anti-rabbit secondary antibody (Invitrogen/Molecular Probes, A21076), as appropriate, and imaged by fluorescence detection.

### Mammalian FLAG ChIP-qPCR

FLAG ChIP-qPCR was used to measure accumulation of Flp-H305L at the FRT locus. HEK-293 and U2OS Flp-In cells were seeded at approximately 5 × 10⁶ cells per 15-cm dish, and TK6 cells were seeded at approximately 2.5 × 10⁶ cells per T25 flask. Cells were treated with doxycycline and other compounds as described above. Unless otherwise indicated, doxycycline was used at 1 µg/mL and MG132 at 30 µM for 8 h. Cells were harvested without formaldehyde crosslinking. Adherent cells were washed twice with ice-cold PBS and collected by scraping. TK6 cells were collected by centrifugation at 800 × g for 3 min and washed with cold PBS. Cell pellets were snap-frozen in liquid nitrogen and stored at −80°C until processing. Frozen pellets were thawed on ice and resuspended in 300 µL RIPA buffer containing 50 mM Tris-HCl, pH 8.0, 150 mM NaCl, 2 mM EDTA, 1% NP-40, 0.5% sodium deoxycholate, and 0.1% SDS, supplemented with protease inhibitors (Thermo Fisher Scientific, A32965). Samples were incubated on ice for 15 min. HEK-293 and U2OS lysates were centrifuged at 5,000 rpm for 10 min, whereas TK6 lysates were centrifuged at 800 × g for 3 min. The supernatant was discarded, and the DNA-containing pellet was gently resuspended in 2.2 mL fresh RIPA buffer.

Chromatin was sheared using a Bioruptor UCD-200 sonicator (Diagenode) for two 15-min cycles with 10-s on/10-s off intervals. Samples were then centrifuged at 800 × g for 3 min at 4°C. A 50-µL aliquot of each cleared lysate was reserved as input, and 2 mL of the remaining lysate was used for immunoprecipitation. Lysates were incubated with 4 µg mouse monoclonal anti-FLAG M2 antibody (Sigma-Aldrich, F3165) for at least 20 h at 4°C with rotation. Protein A/G magnetic beads were then added and incubated for 3 h at room temperature with rotation. Either Pierce Protein A/G Magnetic Beads (Thermo Fisher Scientific, 88803) or Protein A/G Magnetic Beads (MedChemExpress, HY-K0202) were used, with 25 µL bead suspension per immunoprecipitation. Beads were captured on a magnetic rack and sequentially washed for 20 min at room temperature with 1 mL each of TSE150 buffer (20 mM Tris-HCl, pH 8.0, 150 mM NaCl, 2 mM EDTA, 1% Triton X-100, 0.1% SDS), TSE500 buffer (20 mM Tris-HCl, pH 8.0, 500 mM NaCl, 2 mM EDTA, 1% Triton X-100, 0.1% SDS), LiCl buffer (10 mM Tris-HCl, pH 8.0, 250 mM LiCl, 1 mM EDTA, 1% NP-40, 1% sodium deoxycholate), and TE buffer (10 mM Tris-HCl, pH 8.0, 1 mM EDTA). Following the LiCl wash, beads were transferred to fresh tubes. Chromatin was eluted in 100 µL elution buffer containing 1% SDS and 100 mM NaHCO₃ for 30 min at 30°C with rotation. Eluted samples were stored at −20°C until further processing.

For DNA purification, 100 µL PBS was added to each IP eluate to reduce the SDS concentration to 0.5%. Samples were treated with 20 µg RNase A for 1 h at 37°C, followed by 20 µg Proteinase K overnight at 65°C. DNA was purified using the QIAquick Nucleotide Removal Kit (QIAGEN, 28306) with 10 volumes of Buffer PNI and eluted in 60 µL Buffer EB.

Purified ChIP and input DNA were analyzed by SYBR Green qPCR using a QuantStudio 7 Real-Time PCR System (Applied Biosystems). Each 10-µL reaction contained 5 µL PowerUp SYBR Green Master Mix, 0.4 µL primer-pair mix containing 12.5 µM of each primer, 1.6 µL nuclease-free water, and 3 µL DNA template. Reactions were performed in technical duplicate. Cycling conditions were 50°C for 2 min, 95°C for 2 min, followed by 40 cycles of 95°C for 15 s and 60°C for 1 min, followed by melt-curve analysis.

For HEK-293 and U2OS Flp-In cells, the FRT-containing region was amplified using TP9191/TP9192. For TK6 FRT-containing cells, the Upstream, Across, and Downstream regions were amplified using TP10249/TP10459, TP10251/TP10252, and TP10253/TP10254, respectively. Primer sequences are listed in Supp. Table 1.

ChIP enrichment was calculated as percent input from the Ct values of the corresponding input and immunoprecipitated samples, accounting for the input dilution factor. Percent-input values were then normalized to the indicated control condition for each experiment.

### MRE11 ChIP-qPCR

MRE11 ChIP-qPCR in TK6 cells was performed using the same chromatin preparation, sonication, immunoprecipitation, washing, DNA purification, and qPCR procedures described above for mammalian FLAG ChIP, except that cells were formaldehyde-crosslinked before harvest and an anti-MRE11 antibody was used for immunoprecipitation. At the end of treatment, TK6 cultures were agitated at 120 rpm and formaldehyde was added directly to the culture to a final concentration of 1%. Cells were crosslinked for exactly 7 min at room temperature. Crosslinking was quenched with 1.25 M glycine, followed by brief agitation. Cells were then collected by centrifugation, washed with cold PBS, snap-frozen in liquid nitrogen, and stored at −80°C until processing. For immunoprecipitation, 3 µL rabbit anti-MRE11 antibody (Novus Biologicals, NB100-142) was added per sample in place of the anti-FLAG antibody. All subsequent procedures were performed as described above.

### *Saccharomyces cerevisiae* FLAG ChIP-qPCR

Yeast cells were induced with galactose or DAPG and synchronized where indicated as described above. Cells were harvested by centrifugation without crosslinking, washed once with cold PBS, flash-frozen in liquid nitrogen, and stored at −80°C. Cell pellets were resuspended in lysis buffer containing 50 mM HEPES, 140 mM NaCl, 1 mM EDTA, 1% Triton X-100, 0.1% SDS, and protease inhibitors. Cells were disrupted with 0.5-mm zirconia beads by three 45-s cycles of bead beating, with 1-min rests on ice between cycles. Lysates were sonicated using a Bioruptor UCD-200 sonicator (Diagenode) for two 12.5-min cycles at high power with 30-s on/30-s off intervals. Samples were cleared by centrifugation at 13,000 rpm for 15 min at 4°C, and a fraction of each cleared lysate was reserved as input. Cleared lysates were incubated overnight at 4°C with rotation with 5 µg anti-FLAG M2 antibody (Sigma-Aldrich, F1804). Protein A/G magnetic beads (Pierce, Thermo Fisher Scientific, 88803) were then added and incubated for 3 h at 4°C with rotation. Beads were sequentially washed with TSE150, TSE500, LiCl, and TE buffers using the same buffer compositions described above for mammalian FLAG ChIP. DNA–protein complexes were eluted in elution buffer containing 2% SDS at 65°C for 30 min.

Input and immunoprecipitated samples were treated with RNase A and Proteinase K, and DNA was purified using the QIAquick Nucleotide Removal Kit (QIAGEN, 28306). ChIP enrichment was measured by SYBR Green qPCR using a ViiA7 Real-Time PCR System (Applied Biosystems). The Upstream, Across, and Downstream regions were amplified using TP10483/TP10484, TP10460/TP10461, and TP10481/TP10482, respectively (See Supp. Table 1). ChIP signal was normalized to input DNA, and resulting enrichment values were normalized to the indicated control condition for each experiment.

### DNA-associated protein isolation and peptide preparation for mass spectrometry

For the initial mass spectrometry experiment shown in Fig. 5B, WT control TK6 cells lacking Flp-H305L expression were treated with MG132 (30 µM), whereas *TDP1−/−* TK6 cells expressing Flp-H305L were treated with doxycycline (1 µg/mL) and MG132 (30 µM) for 8 h. For the independent mass spectrometry experiment shown in Fig. 5C–G, samples consisted of untreated WT TK6 cells, WT TK6 cells treated with MU147 (25 µM) for 4 d, and MRE11-AID TK6 cells treated with 5-Ph-IAA (10 µM) for 5 d to deplete MRE11. Cells used in the latter experiment did not express Flp-H305L. Immediately before cell harvesting, cell viability was measured using a Bio-Rad TC20 Automated Cell Counter with trypan blue exclusion. For each measurement, 10 µL of cell suspension was mixed with 10 µL trypan blue, and 10 µL of the mixture was loaded onto a counting slide. Three measurements were obtained for each sample and averaged to determine the survival rate used for subsequent normalization of protein abundance. Cell pellets containing approximately 3–5 × 10^6^ TK6 cells per sample were thawed on ice and resuspended in 500 µL Genomic Lysis Buffer from the Quick-DNA Miniprep Kit (Zymo Research). Samples were mixed by pipetting until the cell pellets were completely dispersed and incubated at room temperature for 5–10 min. Lysates were transferred to Zymo-Spin IICR columns and centrifuged at 10,000 × g for 1 min, and the flow-through was discarded. Column-bound material was washed sequentially using two custom wash buffers. Wash buffer 1 (WB1) contained 80% ethanol and 10 mM Tris-HCl, pH 7.4, whereas wash buffer 2 (WB2) contained 75% ethanol, 0.5 M NaCl, and 10 mM Tris-HCl, pH 7.4. Columns were washed twice with 500 µL WB1, once with 300 µL WB2, and twice more with 500 µL WB1. Columns were centrifuged at 10,000 × g for 1 min after each wash. Following the final wash, columns were centrifuged for an additional 3 min to remove residual ethanol and transferred to clean microcentrifuge tubes.

DNA-associated proteins retained on the columns were digested by addition of 100 µL trypsin working solution containing 5 µg/mL Trypsin Gold in 40 mM ammonium bicarbonate. Samples were incubated at 37°C overnight. The following day, columns were centrifuged at 10,000 × g for 1 min to recover the tryptic digest. An additional 100 µL of freshly prepared 20% acetonitrile/2% formic acid was added to each column, followed by incubation for 5 min at room temperature and centrifugation at 10,000 × g for 1 min. The resulting eluates were combined. DNA concentration in the combined Zymo-column eluate was measured using a NanoDrop spectrophotometer before subsequent centrifugal-filter purification. For further peptide purification, 30-kDa molecular-weight-cutoff regenerated-cellulose centrifugal filters (Ultracel YM-30, Millipore) were prewashed with 400 µL 20% acetonitrile/2% formic acid by centrifugation at 14,000 × g for 10 min. The combined eluate was applied to the filter and centrifuged at 14,000 × g for 10 min, and the peptide-containing flow-through was collected. Filters were subsequently rinsed with an additional 50 µL 20% acetonitrile/2% formic acid, and the resulting flow-through was combined with the initial filtrate. Purified peptide samples were dried by SpeedVac without heating and submitted for mass-spectrometric analysis.

### LC-MS/MS and protein identification

Dried peptide samples were submitted to the UT Austin Center for Biomedical Research Support Biological Mass Spectrometry Facility (RRID: SCR_021728) for desalting and LC-MS/MS analysis. Protein identification and quantification were performed using Proteome Discoverer version 2.5.0.400 (Thermo Fisher Scientific). Peak lists were generated using the Spectrum Selector node and searched with Sequest HT against the UniProt canonical reference proteome for *Homo sapiens*, supplemented with the *S. cerevisiae* Flp site-specific recombinase sequence and a contaminant database.

Trypsin was specified as the proteolytic enzyme with up to 2 missed cleavages. Precursor and fragment mass tolerances were set to 10 ppm and 0.6 Da, respectively. Methionine oxidation was set as a dynamic modification. Peptide spectrum matches (PSMs) and protein false discovery rates (FDR) were validated using the Percolator node with a strict and relaxed FDR threshold of 0.01 and 0.05, respectively. Intensity-based label-free quantification was performed on unique peptides using the Minora Feature Detector node and abundances were normalized based on total peptide amount. Total normalized peptide abundance data was further processed in RStudio.

### Proteomic data processing and analysis

Contaminant proteins were removed, and only high-confidence protein identifications with FDR ≤ 0.01 and at least two unique peptides were retained for downstream analysis. Protein abundance values were further normalized to sample input according to: Input-normalized abundance= Total peptide-normalized abundance /[(DNA concentration) × (survival rate)]. Missing values were imputed using values randomly drawn from the distribution of the lowest 5% of abundance values within the corresponding sample group. For each treatment-versus-control comparison, proteins with more than 50% imputed values in the treatment-condition replicates were then excluded. For each protein, mean abundance was calculated for each sample group, and log2 fold change was calculated for treatment relative to the corresponding control. Statistical significance was assessed using Welch’s two-tailed t-test. Benjamini-Hochberg correction was applied separately to each treatment-versus-control comparison to obtain adjusted P values. Volcano plots were generated using log2 fold changes and −log10-transformed unadjusted P values. Selected proteins, including TOP1, were annotated on the plots. See Supp. Table 2.

### TOP1cc immunofluorescence

Asynchronous U2OS cells stably expressing doxycycline-inducible, shMRE11-resistant MRE11-WT or MRE11-H129N were seeded on sterile coverslips and treated with 1 µM doxycycline for 48 h. The corresponding constructs also contained an H1 promoter-driven shMRE11 cassette to deplete endogenous MRE11. Following treatment, cells were washed once with PBS and fixed with 4% paraformaldehyde in PBS for 15 min at 4°C. Cells were washed with PBS and permeabilized with 0.5% SDS in PBS for 5 min. Cells were then washed five times with wash buffer consisting of PBS supplemented with 0.1% Triton X-100 and 0.1% BSA, followed by blocking with 5% BSA in PBS for 1 h at room temperature. TOP1 covalent complexes were detected using a mouse monoclonal anti-TOP1cc antibody (TopoGEN, TG2017-2) diluted 1:400 in 2.5% BSA in PBS for 1 h at room temperature. Following primary-antibody incubation, cells were washed three times with PBS and incubated with goat anti-mouse Alexa Fluor 647 secondary antibody (Thermo Fisher Scientific) at 1:500 together with Hoechst 33342 at 1:3,000 in 1% BSA in PBS for 1 h at room temperature. Cells were subsequently washed three times with PBS and mounted on microscope slides. Fluorescence images were acquired using a Nikon AXR-NSPARC microscope with a 20× objective. TOP1cc fluorescence intensity was quantified using ImageJ and normalized to the corresponding MRE11-WT control.

### Statistical analysis

Data are presented as mean ± SEM unless otherwise indicated. Individual points in bar graphs represent independent biological replicates. Technical replicates, including technical qPCR replicates, were averaged within each biological replicate and were not treated as independent samples for statistical analysis. Statistical analyses were performed in Microsoft Excel.

For comparisons in Figs. 1–4, statistical significance was determined using two-tailed Student’s t tests assuming equal variance. For Fig. 5B and Fig. 5D–G, two-tailed Welch’s t tests assuming unequal variance were used. For Fig. 5J, statistical significance was determined using a two-tailed Student’s t test assuming equal variance using the mean TOP1cc intensities from three independent biological experiments. Significance thresholds were defined as P < 0.05 (*), P < 0.01 (**), P < 0.001 (***), and P < 0.0001 (****); ns indicates not significant. Exact P values are shown for selected non-significant comparisons where indicated.

For proteomic analyses, statistical significance for each protein was assessed using a two-tailed Welch’s t test. Benjamini–Hochberg correction was applied separately to each treatment-versus-control comparison to obtain adjusted P values. Volcano plots display log2 fold changes and −log10-transformed unadjusted P values.

